# RAD51 Paralogs and RAD51 Paralog Complexes BCDX2 and CX3 Interact with BRCA2

**DOI:** 10.1101/2024.10.10.617680

**Authors:** Jacob G. Thrasher, Adeola A. Faunloye, Fabricio S. Justiniano, Kara A. Bernstein, Ryan B. Jensen

## Abstract

Homologous recombination (HR) is an important mechanism for repairing DNA double-strand breaks (DSBs) and preserving genome integrity. Pathogenic mutations in the HR proteins BRCA2 and the RAD51 paralogs predispose individuals to breast, ovarian, pancreatic, and prostate cancer. The RAD51 paralogs: RAD51B, RAD51C, RAD51D, XRCC2, and XRCC3 form two complexes RAD51B-RAD51C-RAD51D-XRCC2 (BCDX2) and RAD51C-XRCC3 (CX3). Similar to BRCA2, loss of RAD51 paralog functions in mammalian cells lead to chromosomal abnormalities, growth defects, disrupted RAD51 foci formation, and PARP inhibitor sensitivity. Despite significant effort over the past three decades, the specific molecular functions of the human RAD51 paralogs have remained elusive due to technical challenges such as low protein expression in human cell lines and instability of the purified proteins. Recent studies have determined the molecular structures of the BCDX2 and CX3 complexes dramatically enhancing our understanding of these challenging proteins. Using multiple approaches, we demonstrate that the RAD51 paralogs interact with BRCA2 at two distinct interaction hubs located in the BRC repeats and the DNA binding domain. We confirm, using a yeast 3-hybrid approach, that human RAD51 paralogs interact directly with BRC repeats one and two (BRC1-2) of BRCA2. Because of the dynamic nature of the RAD51B C-terminal domain (CTD), identified in the recently solved cryo-EM structures, we focused on elucidating the interaction with RAD51B. We determined that BRCA2 interacts with the CTD of RAD51B and not the N-terminal domain (NTD) that is involved in stacking interactions with RAD51C and RAD51D. Furthermore, the interaction with RAD51B is dependent upon an FxxA motif located on a surface exposed region of the CTD. Our study has identified novel interactions between the RAD51 paralogs and BRCA2 and further demonstrated that a previously unrecognized FxxA motif located within a mobile element of RAD51B is critical for the interaction.

## Introduction

The ability to repair DNA lesions from both environmental and endogenous sources is critical for preserving genome integrity and preventing neoplasia[1]. DNA can incur many types of damage, with DNA double-strand breaks (DSBs) being among the most difficult to repair [2]. When DNA DSBs are repaired through mutagenic pathways like non-homologous end-joining (NHEJ) or polymerase theta-mediated end-joining (TMEJ), they can result in chromosomal aberrations, neoplastic transformation, and ultimately contribute to cancer progression and treatment resistance[3, 4]. In contrast, HR is a highly accurate pathway that primarily occurs during S and G_2_ phases and uses a homologous DNA template for error-free DSB repair[5–9]. Defects in either HR or NHEJ can lead to programmed cell death, cellular senescence, genomic scarring, and chromosomal translocations.

During HR, a DNA DSB is resected to produce a 3’ single-stranded DNA (ssDNA) overhang which is rapidly coated by **R**eplication **P**rotein **A** (RPA) to prevent secondary structures, spontaneous annealing, and ssDNA degradation[10, 11]. **BR**east **CA**ncer gene 2 (BRCA2) binds, loads, and stabilizes RAD51 ultimately displacing RPA from the 3’ ssDNA overhang to form a RAD51 nucleoprotein filament[12–14]. The RAD51 nucleoprotein filament is essential for the subsequent strand invasion and homology search steps necessary for HR, but the formation and stabilization of the RAD51 filament is dependent on BRCA2 and other auxiliary factors such as BRCA1, PALB2, and likely the RAD51 paralogs[15–17]. BRCA2 is a key catalyst of HR and pathogenic germline mutations dramatically increase the risk for hereditary breast and ovarian cancers as well as other epithelial tumors [2, 18–22]. BRCA2 has three major domains implicated in different functions of BRCA2: an N-terminal region important for interactions with PALB2, eight BRC repeats involved in RAD51 loading and stabilization onto ssDNA, a DNA binding domain (DBD) composed of an alpha helix region and three tandem oligonucleotide/oligosaccharide-binding folds (OB-folds), and a C-terminal region that includes a C-terminal RAD51-binding (TR2) domain acting as an additional RAD51 binding site (Figure 1A) [23–26]. The TR2 domain of the C-terminal region of BRCA2 contains a highly conserved proline-rich region preceded by a stretch of positively charged residues which are highly conserved. This region, which is largely disordered, interacts with both DMC1, a meiosis-specific recombinase, and RAD51[26, 27]. In addition, the RAD51 paralogs are also implicated in promoting RAD51 filament formation and stabilization, although the molecular mechanisms by which the RAD51 paralogs function in HR requires further investigation[28–32].

**Figure 1.**
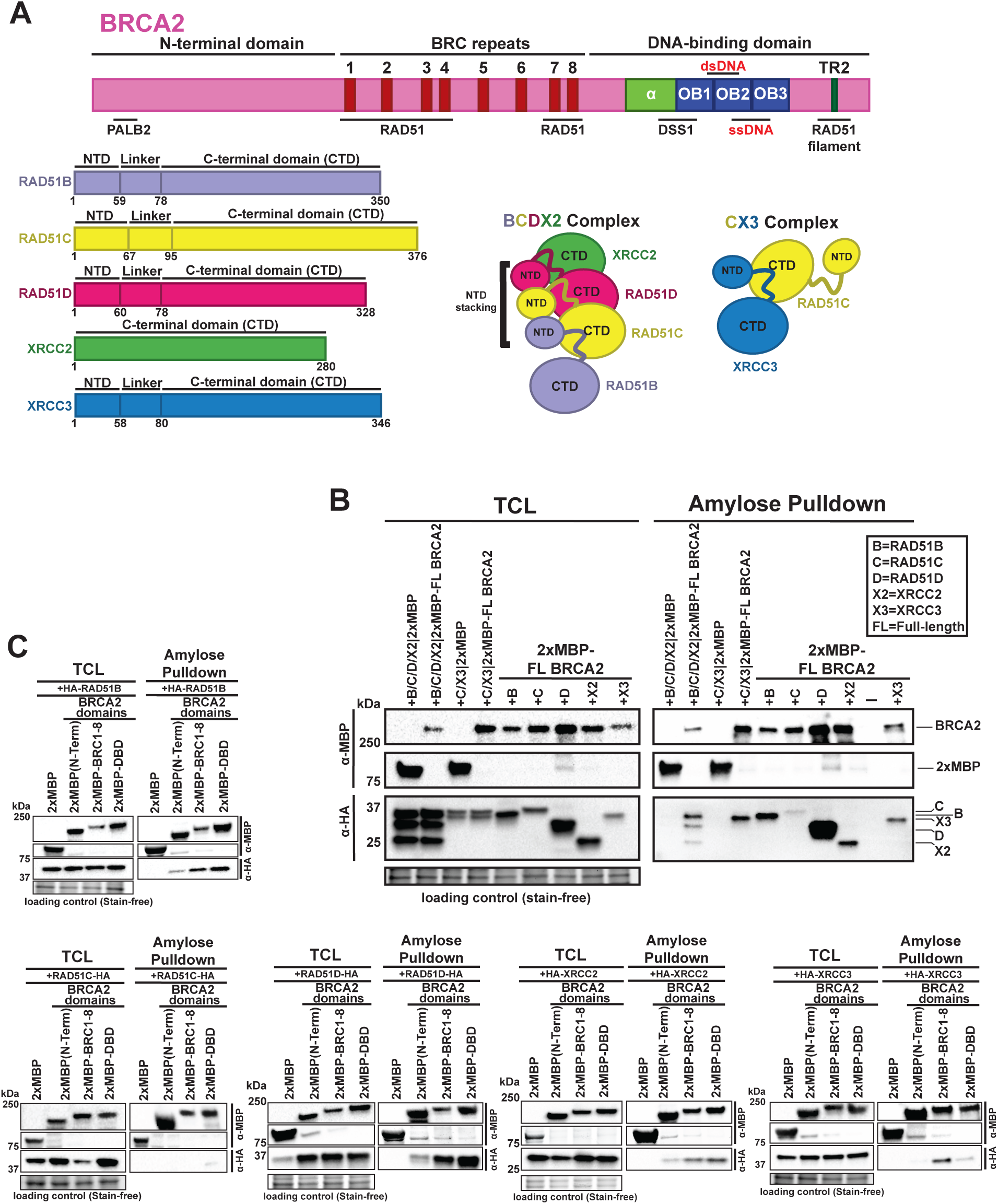
BRCA2 interacts with the RAD51 paralogs and RAD51 paralog complexes. **(A)** Schematic representation of BRCA2 and RAD51 paralogs. BRCA2 (3,418 amino acids) comprises an N-terminal domain, eight BRC repeats (residues 1002–2085), and a C-terminal DNA-binding domain (DBD) with multiple sites binding ssDNA and dsDNA, plus a nuclear localization signal (NLS). RAD51 paralogs contain N-terminal, linker, and C-terminal domains and can form BCDX2 and CX3 complexes as illustrated. **(B)** Full-length BRCA2 interacts with RAD51 paralogs. Western blots of amylose pulldowns showing interactions between 2XMBP-tagged full-length BRCA2 and HA-tagged RAD51 paralogs, both individually and in BCDX2 and CX3 complexes. **(C)** BRCA2 domains interact with individual RAD51 paralogs. Western blots of amylose pulldowns showing interactions between 2XMBP-tagged BRCA2 domains (N-term, BRC1–8, DBD) and HA-tagged RAD51 paralogs. *(Note: Panel C includes five separate pulldown experiments, each with a diGerent RAD51 paralog.)* Stain-free gel images served as loading controls.

In mammalian cells, there are 5 canonical RAD51 paralogs: RAD51B, RAD51C, RAD51D, XRCC2, and XRCC3[33–36]. These RAD51 paralogs share 20-30% sequence identity with RAD51, particularly in the N-terminal domains and conserved Walker A and Walker B motifs which are essential for ATP binding and hydrolysis (Figure 1A) [30, 37]. The RAD51 paralogs have also been shown to form two distinct complexes (Figure 1A): RAD51B-RAD51C-RAD51D-XRCC2 (BCDX2), and RAD51C-XRCC3 (CX3), with RAD51C being the only member of both complexes (Figure 1A) [29, 38–40].

Recent structural studies of BCDX2 revealed how these proteins interact to form the complex[31, 32, 41]. Similar to RAD51, all RAD51 paralogs, except XRCC2, have an N-terminal domain (NTD) and linker domain important for their interactions (Figure 1A) [41]. Unlike RAD51, whose NTD folds back towards its ATPase domain, the NTDs of RAD51C and RAD51D project away from their ATPase domains[31, 32]. In RAD51 presynaptic filaments, the NTDs have few contacts with neighboring subunits, but in the BCDX2 complex, RAD51B, RAD51C, and RAD51D extensively contact NTDs and ATPase domains of neighboring subunits[31, 32, 41]. In the BCDX2 complex, the NTDs of the paralogs play an important role through NTD stacking- a type of interaction not seen with the NTDs of RAD51[31, 32].

The RAD51 paralogs play crucial roles in the DNA damage response and are genetically epistatic to BRCA2 [15]. Knockout of BRCA2 or the RAD51 paralogs results in loss of RAD51 foci in deficient cells and embryonic lethality in RAD51B, RAD51C, RAD51D, XRCC2, and XRCC3 knockout mice [42–48]. RAD51 paralog deficient cells exhibit extreme sensitivity to cross-linking agents and mild sensitivity to ionizing radiation [35, 44, 49, 50]. Mutations in HR proteins that lead to HR deficiency significantly elevate the risk for various cancers by compromising genome integrity[19]. Inherited mutations in both BRCA2 and the RAD51 paralogs are associated with a higher risk of breast and ovarian cancer, however, because of HR deficiency due to loss-of-heterozygosity within the tumor, these patients often respond favorably to PARP inhibitors [32, 41, 51–57]. Despite many parallels, direct interaction between BRCA2 and the RAD51 paralogs has not been characterized to date.

In this study, we used multiple approaches to demonstrate protein-protein interactions between the RAD51 paralogs and BRCA2 [13, 18]. Interactions were detected between the BRC repeats region (BRC1-8) of BRCA2 and all RAD51 paralogs except for RAD51C. We also detected interactions between the DNA binding domain (DBD) of BRCA2 and the RAD51 paralogs. Using a yeast 3-hybrid approach, we determined multiple direct interactions between BRC1-2 and multiple RAD51 paralogs in the BCDX2 and CX3 complex. Considering the recent cryo-EM studies describing the mobility of the CTD domain of RAD51B relative to the core of the BCDX2 complex[31, 32], we further explored the specific interactions of RAD51B with BRCA2. Because of the interactions with the BRC repeat region of BRCA2, upon close visual inspection of the sequences, we identified FxxA motifs in four of the five RAD51 paralogs. Using AlphaFold predicted structures and recent cryo-EM structures of the BCDX2 complex, we found that the FxxA motif of RAD51B is solvent-exposed, suggesting a potential interaction hub for RAD51B with other HR proteins such as BRCA2 and RAD51. We created a RAD51B F78A mutant to disrupt this motif and determined its impact on interactions with BRCA2 and the other RAD51 paralogs. To better understand the physiological impact of the RAD51B NTD and CTD domains and the RAD51B F78A mutant, we expressed the proteins in a RAD51B knockout cell line and performed RAD51 foci analysis following DNA damage.

## Results

### BRCA2 interacts with individual RAD51 paralogs and the RAD51 paralog complexes: BCDX2 and CX3

Disrupting BRCA2-RAD51 interactions either through genetic mutations or chemical inhibitors result in HR deficiency in cells [18, 58]. Considering the importance of the BRCA2-RAD51 interaction and the shared evolutionary origin and structural similarities of RAD51 and the RAD51 paralogs, we speculated the RAD51 paralogs could directly interact with BRCA2 to regulate RAD51 functions. This regulation is needed because RAD51 activity must be tightly controlled to maintain genomic stability. Unregulated or misregulated RAD51 can lead to inappropriate recombination events, chromosomal rearrangements, or genomic instability, all of which are hallmarks of cancer[59, 60]. The interaction between RAD51 paralogs and BRCA2 could ensure that RAD51 filament formation happens in a coordinated and efficient manner. To determine whether the RAD51 paralogs interact with the full-length BRCA2 protein, we first tested whether recombinant expression in human 293T cells would identify specific interactions. A tandem repeat of the maltose binding protein tag (designated 2XMBP) fused to the N-terminus of BRCA2 overcomes the limited expression issues associated with the untagged recombinant BRCA2 protein. We co-expressed both 2XMBP-BRCA2 and HA-tagged RAD51 paralogs either individually or together as the two complexes: BCDX2 or CX3 (Figure 1B). Taking advantage of the 2XMBP tag on BRCA2, we performed amylose pulldowns to capture the protein complexes (Figure 1B). As BRCA2 and the RAD51 paralogs are DNA binding proteins, all cellular lysates were treated with Benzonase nuclease to degrade all forms of DNA and RNA that could potentially bridge the proteins in the amylose pulldown experiments.

Strikingly, the full-length BRCA2 protein pulled down a significant amount of RAD51D (Figure 1B) followed by RAD51B and lesser amounts of RAD51B, XRCC2, XRCC3 (Figure 1B). With respect to the two paralog complexes, RAD51B, RAD51D, and XRCC2 were pulled down from the BCDX2 complex (Figure 1B) and XRCC3 from the CX3 complex (Figure 1B), however, RAD51C was notably absent and/or scarcely detectable in the individual and complex pulldowns. Non-transfected cells were used as a control for any non-specific bands and the 2XMBP tag alone was transfected with both RAD51 paralog complexes (Figure 1B) to further control for any non-specific binding of the paralogs to the 2XMBP tag or the amylose beads. Given the known interactions between the paralogs, it was somewhat unexpected that the one component common to both complexes, RAD51C, displayed such a weak interaction signal. While it is unlikely that endogenous RAD51 paralogs are bridging the interactions between the recombinantly expressed BRCA2 and other members of the BCDX2 and CX3 complex, the possibility still exists that other proteins in the cellular extract could mediate indirect interactions, and thus, we explored the interactions in a heterologous expression system as described below.

### The RAD51 paralogs interact directly with the BRC repeats of BRCA2

We wanted to determine if a specific region of BRCA2 could be mapped to the interaction site for the RAD51 paralogs to better understand how these interactions may impact RAD51 paralog functions depending on with which domains of BRCA2 the RAD51 paralogs interact. We divided BRCA2 into three fragments fused to the 2XMBP tag: N-terminal domain (2XMBP-N-TERM), BRC1-8 repeats (2XMBP-BRC1-8), and a construct comprising the DNA-binding and C-terminal domain (2XMBP-DBD) (Figure 1A). Through these amylose pulldowns we observed intense signal for both RAD51B and RAD51D pulled down by both the BRC1-8 and DBD of BRCA2 (Figure 1C), and weak signal for RAD51B and RAD51D being pulled down by the N-term of BRCA2 (Figure 1C). No interactions were detected between RAD51C and any of the BRCA2 domains for XRCC2, interactions were detected for BRC1-8 and DBD, but the signal was weaker than that for RAD51B or RAD51D (Figure 1C). Very faint signal was detected for XRCC2 interaction with BRCA2 N-term (Figure 1C). For XRCC3, an interaction was detected for BRC1-8, and a very faint signal was detected for DBD (Figure 1C). No interaction was detected between XRCC3 and the N-term of BRCA2 (Figure 1C).

The interaction between the BRC repeats and the RAD51 paralogs were particularly compelling as the BRC repeats function as an interaction hub for RAD51 and are critical for RAD51 loading and filament formation[13, 14, 61, 62]. We divided the BRC repeats of BRCA2 into the following fragments fused to the 2XMBP tag: BRC1-4, BRC5-8, BRC1-2, BRC3-4, BRC5-6, BRC7-8 and performed amylose pulldowns with the same HA-tagged paralog constructs to observe with which BRC repeats the RAD51 paralogs are interacting (Figure 2A). We see interaction between BRC1-4 and RAD51B, RAD51D, XRCC2, and XRCC3 with the strongest interactors indicated by the highest signal in the pulldowns being RAD51B and XRCC3 (Figure 2A). We also observe interactions between BRC5-8 and RAD51B, RAD51D, XRCC2, and XRCC3, though the signal for these interactions is consistently weaker than the interaction between BRC1-4 and the RAD51 paralogs (Figure 2A). For the BRC doublets, the strongest signal for interaction is between BRC1-2 and the RAD51 paralogs, while there are no readily detectable interactions between BRC3-4 and any of the RAD51 paralogs (Figure 2A). While a signal for interaction between BRC5-6 and BRC7-8 with RAD51B, RAD51D, XRCC2, and XRCC3 is observed, this signal is weaker than that of the interactions with BRC1-2 (Figure 2A). Since BRC1-4 and BRC1-2 are important for nucleating the RAD51 filament, the interactions detected at these BRC repeats with the RAD51 paralogs could be important for RAD51 filament nucleation.

**Figure 2.**
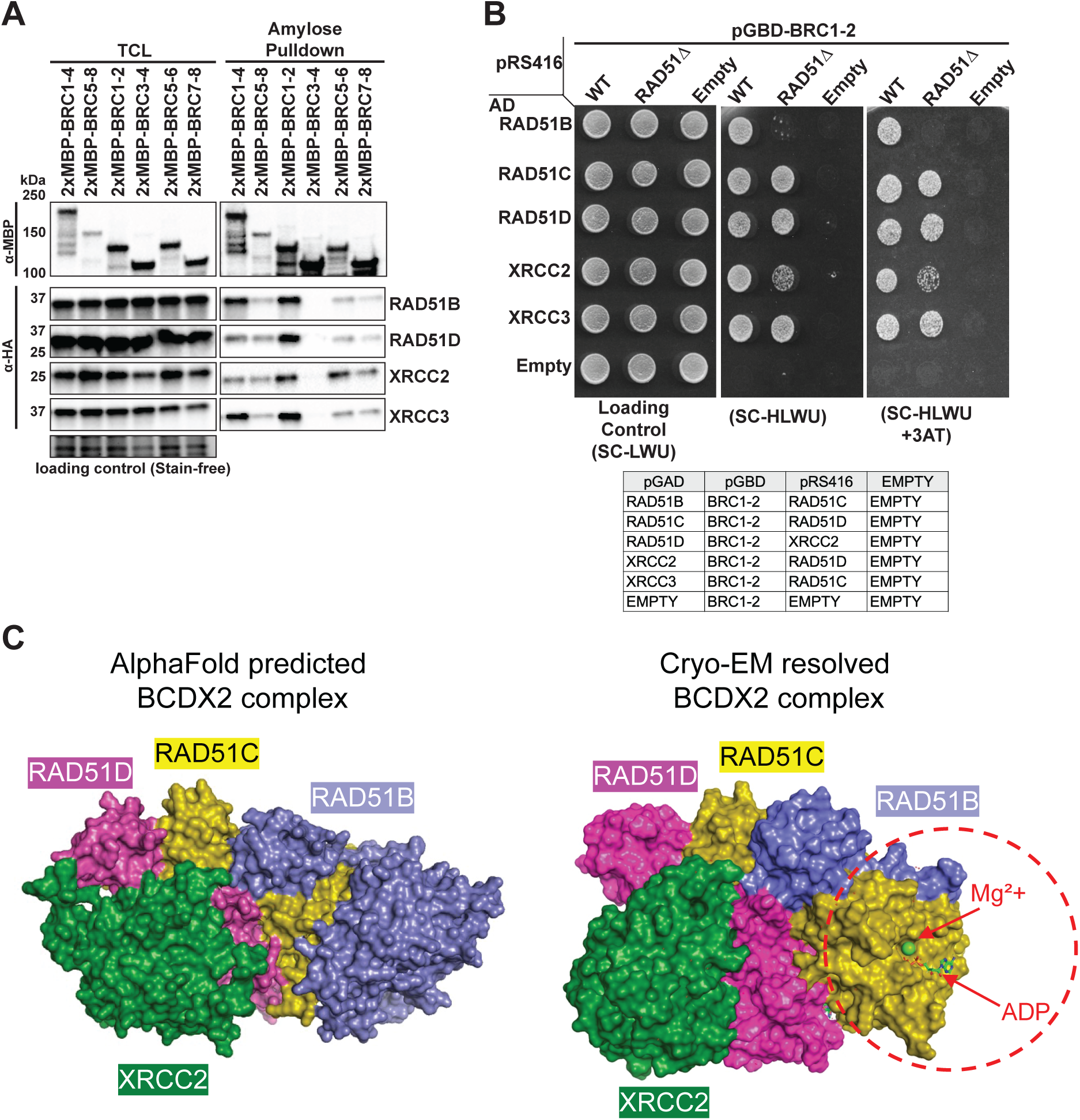
RAD51 paralogs interact with BRCA2 BRC repeats, directly with BRC1-2. **(A)** RAD51 paralogs interact with specific BRC repeats. Western blots of amylose pulldowns from cells co-expressing 2XMBP-tagged BRC repeats (BRC1–4, BRC5–8, BRC1–2, BRC3–4, BRC5–6, BRC7–8) and HA-tagged RAD51 paralogs, showing interactions between RAD51 paralogs and specific BRC repeats. Stain-free gel images served as loading controls. The top anti-MBP and bottom stain-free panels are representative; each anti-HA panel corresponds to a separate experiment for each paralog. **(B)** BRC1–2 repeats exhibit Y3H interaction with BCDX2 and CX3 members in RAD51 knockout background. Yeast-three-hybrid assays in RAD51 KO strains expressing GAL4 BD–BRC1–2 and GAL4 AD–RAD51 paralogs, along with a stabilizing pRS416 vector. Equal cell numbers were spotted on control and selection plates. Growth after 2 days at 30°C indicates protein–protein interaction. Experiments were performed in triplicate; representative images are shown. **(C)** AlphaFold-predicted model (ModelArchive: ma-a54ps) and Cryo-EM structure (PDB: 8OUZ) of the BCDX2 complex. The tetrameric complex consists of RAD51B (purple), RAD51C (yellow), RAD51D (pink), and XRCC2 (green). In both models, N-terminal domain (NTD) stacking stabilizes the complex; XRCC2 interacts with NTDs of RAD51C and RAD51D and the C-terminal domain (CTD) of RAD51D. RAD51B, RAD51C, and RAD51D have outward-projecting NTDs forming trans interactions crucial for stability. Due to RAD51B CTD dynamics, a truncated RAD51B was used in the Cryo-EM structure (missing portion indicated by a dotted line); the AlphaFold model includes full-length RAD51B. The complex requires ATP/Mg²⁺ for stability; nucleotide binding was observed in ATP-binding sites of RAD51C, RAD51D, and XRCC2, with ADP modeled into RAD51C’s site in the Cryo-EM structure.

Considering the inability to rule out bridging proteins in the amylose pulldown assays from human cell extracts, we wanted to determine if the interactions between the paralogs and the BRC repeats were direct using a heterologous expression system. We used a yeast three-hybrid (Y3H) approach, rather than two-hybrid, as prior studies demonstrated a requirement for co-expression of human RAD51 paralogs in yeast as sub-complexes (e.g. BC, CD, DX2) to ensure robust expression and RAD51 paralog stability [41]. Based on the interactions observed in our amylose pulldown assays, we focused on the BRC1-2 region of BRCA2. We co-expressed BRC1-2 fused to the GAL4 DNA binding domain as the bait, each RAD51 paralog fused to the activation domain of GAL4 as the prey, and its cognate binding partner, untagged, in the pRS416 expression vector (Supplementary Figure 1). We performed the Y3H experiment using increasingly stringent conditions to determine the relative strength of each protein-protein interaction (Supplementary Figure 1). Upon analysis of the Y3H results, all the canonical RAD51 paralogs exhibited a Y3H interaction with BRC1-2 where RAD51C was the weakest interactor, a trend that tracked with our amylose pull-down approach using recombinant protein expression in human 293Ts (Figure 1B). SWSAP1, a non-canonical RAD51 paralog and member of the Shu complex, showed no Y3H interaction with BRC1-2 (Supplementary Figure 1). RAD51 was used as a positive control and displayed the strongest interaction (Supplementary Figure 1). Considering the weaker interaction on the most stringent condition in the Y3H experiments and the inability of RAD51C to be significantly pulled down with full-length BRCA2 or any BRCA2 domain in the amylose pulldown experiments (Figures 1B & 1C), it is possible that RAD51C does not directly interact with BRCA2. Because the Y3H approach requires co-expression of the RAD51 paralog sub-complexes, the weak interaction we observe between RAD51C and BRC1-2 could be mediated by the cognate binding partner RAD51D which shows growth for each Y3H experiment it is expressed in regardless of expression via pGAD or pRS416. XRCC3, a member of the CX3 complex, shows interaction even under stringent conditions, suggesting the CX3 complex, too, interacts directly with BRC1-2 likely through interactions with XRCC3 (Supplementary Figure 1). These Y3H interactions provide compelling evidence that both RAD51 paralog complexes BCDX2 and CX3 directly interact with the BRC1-2 region of BRCA2.

Previous work on BRCA2 has shown that human full-length BRCA2 can, in fact, bind to yeast Rad51[13]. Considering the high homology between the two orthologs, being ∼67% identical and the conservation of BRCA2-RAD51 interaction throughout evolutionary history, human BRCA2 interacting with yeast Rad51 is unsurprising. The interaction between the human RAD51 paralogs and human RAD51 has been shown through loss of RAD51 foci formation in RAD51 paralog knock out cells, but the specific sites of interaction between the human BCDX2 complex and RAD51 is still unresolved [15, 29, 63]. Recent surface plasmon resonance (SPR) experiments, fluorescence anisotropy experiments, and FRET experiments all provide compelling evidence that RAD51B, rather than the other BCDX2 paralogs, is responsible for the BCDX2 interaction with RAD51, but those experiments were unable to clearly distinguish if RAD51B was interacting with RAD51, ssDNA, or both [31, 32]. To date, it seems there is no experimental evidence supporting or refuting any of the RAD51 paralogs’ ability to interact with yeast Rad51. Even without this evidence, though, we performed the Y3H experiments in a *RAD51* knockout *S. cerevisiae* competent PJ69-4α yeast strain (*rad51Δ*) to determine if the endogenous yeast Rad51 was bridging any of the Y3H interaction observed in our experiments (Figure 2B). Surprisingly, in the *rad51Δ* Y3H, the interaction between BRC1-2 and RAD51B is lost even on the least stringent plate (SC-HLWU), and interaction between BRC1-2 and XRCC2 is weaker in the *rad51Δ* Y3H compared to the WT Y3H (Figure 2B). Although these results determine that RAD51B does not interact directly with BRC1-2, these results do not preclude RAD51B from interacting directly with other regions of BRCA2 such as other BRC repeats or DBD. The loss of RAD51B-BRC1-2 in the *rad51Δ* background, though, does imply that RAD51B does interact with yeast Rad51 (Figure 2B). Like with BRCA2, human RAD51B interaction with yeast Rad51 is strong evidence that RAD51B-RAD51 interactions are conserved, which strongly supports previous evidence that RAD51B interacts with RAD51 to coordinate RAD51-BCDX2 interaction.

Considering RAD51C was expressed by pRS416 along with RAD51B in these experiments, and the co-expressed RAD51C did not preserve this interaction with BRC1-2 in the absence of Rad51, we believe that RAD51C does not directly interact with BRC1-2 either. The growth seen in the WT Y3H and Rad51Δ Y3H with pAD-RAD51C are likely being bridged by direct interaction of BRC1-2 with the RAD51D that is co-expressed by pRS416 in these experiments (Figure 2B). Taken along with the lack of interaction between BRCA2 and RAD51C in the amylose pulldown experiments, we think it is likely that RAD51C, as a member of either BCDX2 or CX3, does not directly interact with BRCA2, and rather the direct interactions of those complexes and BRCA2 are mediated by other RAD51 paralogs in those respective complexes.

The reduction in growth seen for BRC1-2 XRCC2 interaction in Rad51Δ Y3H compared to the WT Y3H suggest that while BRC1-2 and XRCC2 do interact directly, the interactions between BRC1-2 and XRCC2 could be stabilized by the presence of Rad51(Figure 2B). In every Y3H experiments, both WT and Rad51Δ, we see strong signal for interaction between BRC1-2 and RAD51D; therefore, we are confident there is strong direct interaction between RAD51 and BRCA2 at BRC1-2.

### The C-terminal domain of RAD51B bridges the interaction between RAD51B and BRCA2

Two recent cryo-EM analyses of the BCDX2 complex have provided immense detail of the composition of the complex, but both structures required deletion of the RAD51B C-terminal domain (CTD) due to its dynamic nature (Figure 2C)[31, 32]. Another recent study detailing various RAD51C cancer variants, many of which disrupt BCDX2 complex formation, utilized AlphaFold to predict the structure of the BCDX2 complex containing the full-length RAD51B (Figure 2C) [41]. It was also determined, using purified RAD51B-RAD51C (BC subcomplex) and purified RAD51D-XRCC2 (DX2 subcomplex), that while all RAD51 paralogs have the capacity to bind ATP, the ATP hydrolytic core of BCDX2 resides within the BC subcomplex, with the DNA-stimulated ATPase activity or RAD51B and RAD51C being coupled [31, 32].

The mobile nature of RAD51B in the BCDX2 complex motivated us to determine if the interactions we saw between BRCA2 and RAD51B in the amylose pulldowns (Figure 1C) were mediated by the NTD, the CTD, or both. To determine this, we created two constructs designed to express the N-terminus of RAD51B (RAD51B(NTD); a.a. 1-67) or the C-terminus (RAD51B(CTD); a.a. 67-350) (Figure 3A). We first characterized the RAD51B domain interactions with the BCDX2 complex itself in our amylose pulldown system. Surprisingly, we found that RAD51B(CTD), like full-length RAD51B, can be pulled down with 2XMBP-RAD51C alone (Figure 3B), but RAD51B(NTD) requires co-expression with RAD51D and XRCC2 (Figure 3B & Supplementary Figure 2A). These data suggest the interaction between RAD51B(CTD) and RAD51C are sufficient to persist (without the NTD of RAD51B), but that N-terminal stacking between all members of the BCDX2 complex are required to stabilize the interaction between RAD51B (NTD) and RAD51C (Supplementary Figure 2A).

**Figure 3.**
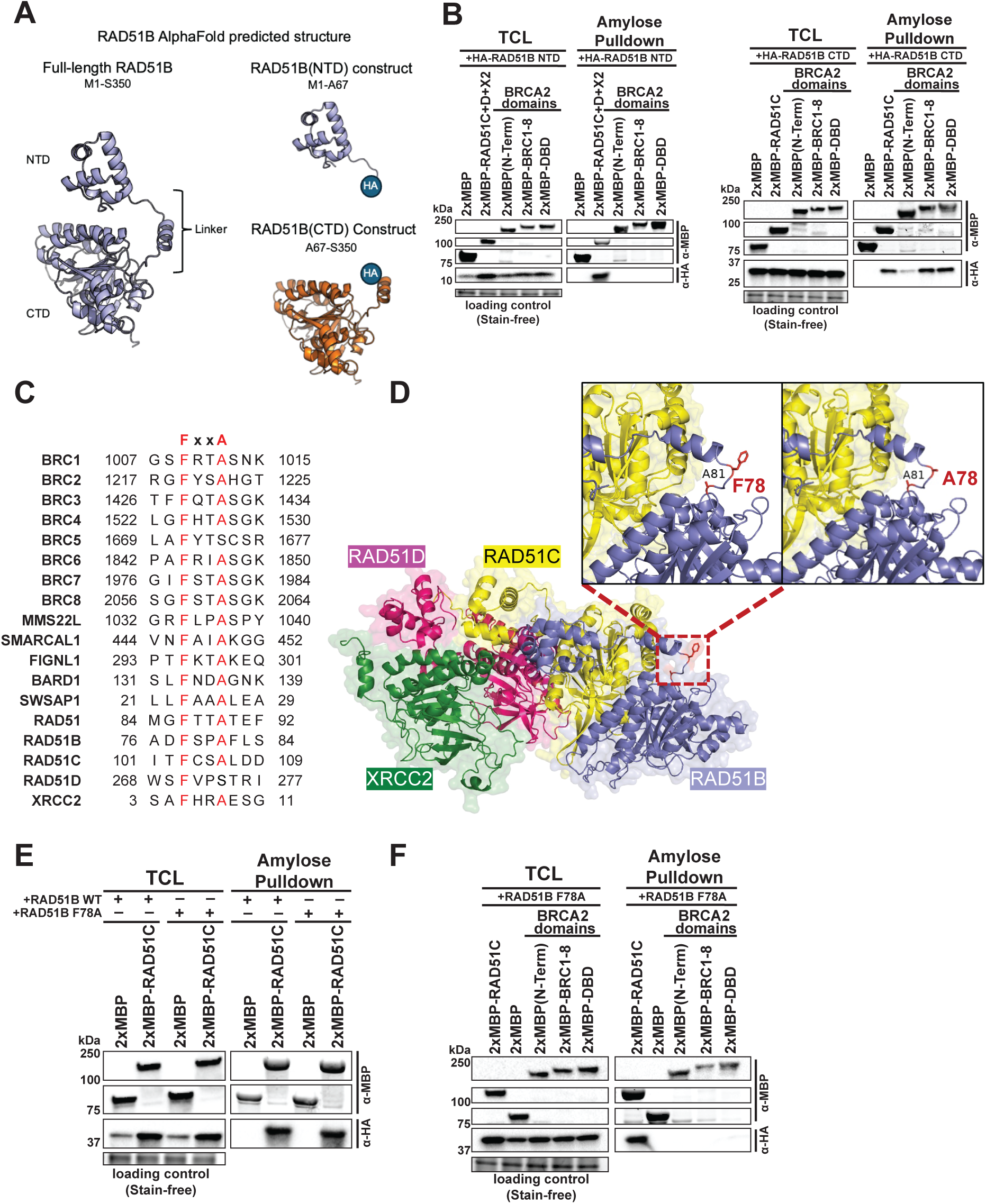
FxxA motif present in RAD51 paralogs, RAD51B FxxA motif critical for BRCA2 interactions. **(A)** AlphaFold-predicted model of RAD51B. Left: Full-length RAD51B with N-terminal domain (NTD), linker domain, and C-terminal domain (CTD) labeled. Right, top: RAD51B NTD with a C-terminal HA tag used to probe NTD-mediated interactions. Right, bottom: RAD51B CTD with an N-terminal HA tag used to investigate CTD-mediated interactions. **(B)** RAD51B domains interact di]erently with BRCA2 and RAD51C. Left: RAD51B NTD interacts with RAD51C (when co-expressed with RAD51D and XRCC2) but not with BRCA2 domains (N-term, BRC1–8, DBD). Right: RAD51B CTD interacts with RAD51C, BRC1–8, and DBD, with a weak interaction with BRCA2 N-term. Western blots of amylose pulldowns show these interactions. **(C)** Multiple sequence alignment (MSA) of FxxA motif across BRCA2 BRC repeats, RAD51 paralogs, and other DNA repair proteins. Key residues are highlighted in red. RAD51D contains an FxxS motif, similar to BRC5 of BRCA2. **(D)** AlphaFold-predicted BCDX2 structure highlighting RAD51B FxxA motif. Ribbon diagram showing RAD51B (purple), RAD51C (yellow), RAD51D (pink), and XRCC2 (green). RAD51B residues F78 and A81 (red) are emphasized. **(E)** RAD51B F78A mutant interacts with RAD51C similarly to wild type. Western blots of amylose pulldowns showing interactions between 2XMBP-RAD51C and HA-tagged RAD51B WT or F78A mutant. **(F)** F78A mutation disrupts RAD51B–BRCA2 interactions. Western blots of amylose pulldowns showing that RAD51B F78A is pulled down by RAD51C but not by BRCA2 domains.

To further refine the interactions between BRCA2 and the RAD51B NTD and CTD domains, we performed amylose pulldowns breaking apart the full-length BRCA2 protein into three regions: the N-terminus (N-TERM), the BRC repeats (BRC1-8), and the DNA binding domain (DBD). None of the BRCA2 fragments were able to pull down the NTD of RAD51B (Figure 3B). In contrast, the CTD of RAD51B was pulled down by both BRC1-8 and the DBD of BRCA2 with a similar signal intensity to the interaction with RAD51C (Figure 3B). The RAD51B CTD interaction with the BRCA2 N-terminus was considerably weaker than BRC1-8 and DBD (Figure 3B). The data suggest that the CTD of RAD51B plays an important role in the interaction with BRCA2 while, consistent with the Cryo-EM and AlphaFold predicted structures, the NTD anchors RAD51B to CDX2 in the BCDX2 complex.

### The FxxA motif in RAD51B is necessary for binding to BRCA2

Interactions between the BRCA2 BRC repeats and RAD51 are mediated by FxxA motifs (Supplementary Figure 2B), a binding motif that has recently been implicated in many protein interactions involved in HDR and fork protection [64, 65]. This motif is comprised of a conserved phenylalanine, two variable amino acids, and a conserved alanine. A crystal structure of the BRC4 peptide in complex with the catalytic domain of RAD51 revealed that the FxxA motif of BRC4 engages hydrophobic binding pockets on the RAD51 surface through the phenylalanine (F1524) and alanine (A1527) (Supplementary Figure 2B)[58, 66]. The RAD51 FxxA motif is not only critical for BRC-RAD51 interactions, but also implicated in RAD51 oligomerization through FxxA binding to the catalytic domain of a cognate RAD51[66–68].

Although a structure-based sequence alignment of the RAD51 paralogs performed on the cryo-EM BCDX2 structure determined that the FxxA motif was not conserved at the same location as in RAD51[31], we visually inspected the amino acid sequences of the RAD51 paralogs and found FxxA motifs in RAD51B, RAD51C, and XRCC2 (Figure 3C). An FxxS sequence, found in the BRC5 repeat of BRCA2 in lieu of FxxA, was identified in RAD51D (Figure 3C). The only RAD51 paralog lacking FxxA and FxxS sequences was XRCC3. It is likely these sequences were not identified in the structure-based sequence alignments due to their distal locations from the motif site in RAD51. Considering the FxxA/FxxS motifs can be found at various positions in BRCA2, BARD1, ZRANB3, MMS22L, RECQL5, SWSAP1, FIGNL1, and other proteins, we were motivated to further explore their importance (Figure 3C) [14, 23, 64, 69–73].

To determine which RAD51 paralog FxxA motifs could be important for BRCA2 interactions, we examined the sequence locations in the recent cryo-EM structures and generated an AlphaFold predicted structure of the BCDX2 complex. The FxxA residues in RAD51B and XRCC2 are not resolved in the cryo-EM structures but are resolved for RAD51C and RAD51D[31, 32]. Both F103 and A106 in RAD51C are buried and are not solvent exposed so it is not likely those residues are involved in BRCA2 interactions (Supplementary Figure 3A). Although a patient mutation, RAD51C(F103V), was recently characterized and found to be HR deficient and unable to interact with RAD51B, RAD51D, and XRCC3 [41]. In the cryo-EM BCDX2 complex structure, the FxxS sequence found in RAD51D sits at the interface between RAD51D and XRCC2 nearby the ATP molecule at this interface (Supplementary Figure 3B) [31, 32]. The aromatic ring of RAD51D F270 is in close contact with the phosphate tail of the ATP molecule located between the RAD51D-XRCC2 interface (Supp Figure 3B). Although the FxxS sequence in RAD51D seems to be important for the DX2 interaction, we did not deem that particular motif as a good candidate for a site of direct interaction between RAD51D and BRCA2 considering it is not surface exposed in the BCDX2 complex (Supp Figure 3B). Using the AlphaFold predicted model of BCDX2, we found that XRCC2 F05 is in proximity to the NTD of RAD51D, and while XRCC2 F05 appears surface exposed in the published AlphaFold predicted model, the pLDDT values were not published with the BCDX2 AlphaFold predicted structure. We regenerated the BCDX2 complex with AlphaFold3, and the average pLDDT for XRCC2 F05 is 27.68 which suggest very low confidence in this position (Supplementary Figure 3B). Considering the proximity of XRCC2 F05 and the NTD of RAD51D and the low pLDDT value, it is unclear is XRCC2 F05 is surface expose or interacting with residues of RAD51D. In the FxxA motif of RAD51B, on the other hand, both the F78 and A81 residues are solvent exposed, are not occupied by intra-complex interactions, and are located within the linker region between the NTD and CTD (Supplementary Figure 3B). Considering the more favorable location and potential for inter-molecular contacts with other proteins, we decided to pursue the FxxA motif in RAD51B, though we think the FxxA motif in XRCC2 warrants further exploration in future experiments.

The FxxA motif in RAD51B contains residues F78-A81 and is located within the CTD construct used in earlier experiments (see Figure 3A). Additionally, since interactions with BRCA2 were observed with the CTD of RAD51B (Figure 3B), we speculated that the FxxA motif is important to mediate the interaction. We used site-directed mutagenesis to generate an F78A mutation in the FxxA motif to determine if RAD51B with a compromised FxxA could still interact with its BCDX2 binding partner, RAD51C, and then BRCA2. Amylose pulldowns were performed from the recombinantly expressed proteins in 293T cells using 2XMBP-RAD51C as the bait and either wild-type RAD51B or the F78A mutant as prey (Figure 3E). Both wild type and the RAD51B F78A mutant were pulled down at similar levels by RAD51C (Figure 3E). While full-length wild type RAD51B and RAD51(CTD) were strongly pulled down with the BRC1-8 and DBD regions of BRCA2, with little signal with the BRCA2 N-terminal construct (Figure 3B), none of the BRCA2 domains pulled down RAD51B F78A (Figure 3F). The loss of F78 in RAD51B abrogates the interaction between any BRCA2 domain and RAD51B F78A but does not affect RAD51B-RAD51C interaction (Figures 3E & 3F), suggesting the FxxA motif in RAD51B is critical for interaction with BRCA2. Considering the results from the Y3H and Rad51Δ Y3H (Figure 2B & Supplementary Figure 1A), it is unclear if RAD51B F78A can no longer bind with BRCA2 because of a loss of direct interaction, or if the BRCA2-RAD51B interactions are mediated by RAD51 and the loss of the FxxA motif in RAD51B impacts its binding with RAD51 which, in turn, impacts the RAD51-mediated BRCA2-RAD51B interaction. Better understanding how the F78A mutation impacts RAD51B-RAD51 binding will help contextualize these results.

### Full-length RAD51B is required for RAD51 foci formation

Nuclear RAD51 foci formation in response to ionizing radiation (IR) damage is an established biomarker for DSB repair mediated by HR [74, 75]. Previous studies have shown that individual RAD51 paralog knockout cell lines do not form RAD51 foci in response to IR, but RAD51 foci formation can be rescued by complementation with their respective wild-type proteins [63]. To further characterize RAD51B and the FxxA motif, we generated stable cell lines by complementing U2OS RAD51B knockout (BKO) cells with wild-type RAD51B, RAD51B(NTD), RAD51B(CTD), and RAD51B F78A (Figure 1C0). Without IR treatment, low rates of spontaneous RAD51 foci formation were evident: BKO (1.5%), RAD51B-WT (7.9%), RAD51B(NTD) (4.9%), RAD51B(CTD) (2.2%), and RAD51B F78A (6.0%) (Figure 4A & 4B). Four hours post 4Gy IR treatment, 55.5% of wild-type RAD51B complemented cells were positive for RAD51 foci (defined as ≤5 foci/nucleus) compared to a 10.9% positive rate for RAD51B knockout cells (Figure 4A & 4B). Neither RAD51B(NTD) nor RAD51B(CTD) complemented cells could rescue with their RAD51 foci positivity rates being 9.5% and 10.1% respectively (Figure 4A & 4B). The cells expressing RAD51B F78A exhibited an intermediate phenotype, partially rescuing RAD51 foci formation with 39.1% of cells positive for RAD51 foci. Together, the data indicate that both the NTD and CTD of RAD51B are necessary for proper RAD51 foci formation in response to IR and that the RAD51B F71A mutation, located within a putative FxxA motif, moderately reduces the efficiency of RAD51 foci formation.

**Figure 4.**
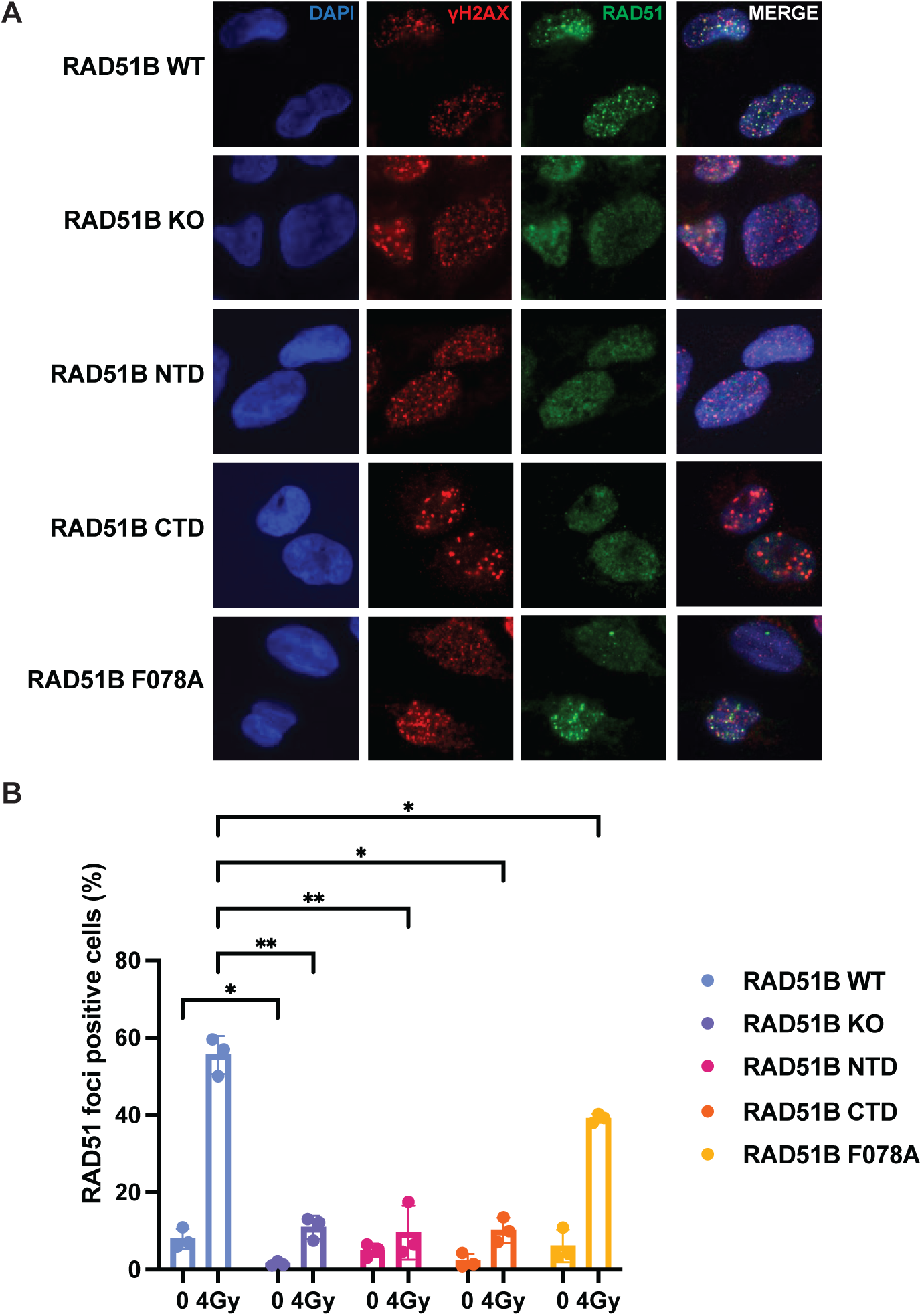
IR-induced RAD51 foci in RAD51B KO, truncation, and mutant stable U2OS cell lines. **(A)** Representative immunofluorescence images of U2OS-derived RAD51B knockout cells (BKO) and BKO cells stably complemented with RAD51B full-length wild type (RAD51B WT), RAD51B N-terminal domain (RAD51B NTD), or RAD51B C-terminal domain (RAD51B CTD), 4 hours post-irradiation (4 Gy). Nuclei are stained with DAPI (blue); γH2AX foci are shown in red; RAD51 foci are shown in green. **(B)** Quantification of the percentage of RAD51 foci-positive cells (≥5 RAD51 foci per nucleus) 4 hours post-treatment with or without IR (4 Gy). Data represent mean ± SEM of three independent experiments. Statistical significance is denoted as (*) p < 0.05 and (**) p < 0.01.

## Discussion

In this work, we demonstrate the novel finding that BRCA2 interacts with the RAD51 paralogs through direct protein-protein interactions. Our study highlights the concept that these tumor suppressor proteins, critical for HR, work closely together to stabilize and enforce the formation of the RAD51 filament. We also provide multiple lines of evidence that BRCA2 interacts with the BCDX2 and CX3 complex in both human cells and in a heterologous yeast expression system. We find that BRCA2 interacts with RAD51B, RAD51D, XRCC2, and XRCC3, yet, surprisingly, only minimally interacts with RAD51C, the one component common to both protein complexes. We then localized the sites of interaction on BRCA2 to the BRC repeats and the DBD. As the BRC repeats are a major hub for RAD51 interactions and regulation, we focused our studies on this domain. One weakness of the amylose pull-down system is that proteins are co-expressed and captured from the same cells so it is possible that endogenous proteins could bridge these interactions. However, it is notable that the lack of RAD51C binding in this approach suggests the endogenous RAD51 paralogs present in the cell extracts are not mediating the detected interactions. Nonetheless, it is still possible that other endogenous proteins could bridge the interactions we observed. To overcome this limitation, we used a heterologous yeast 3-hybrid system to confirm the RAD51 paralogs interact directly with BRCA2.

The full-length BRCA2 protein at 384kDa, has proven difficult to incorporate into yeast 2-hybrid (Y2H) and Y3H experiments, but previous work using smaller fragments and domains of BRCA2 have been successful. Mutations within the BRC repeats can lead to HR deficiency and increase an individual’s predisposition to cancer. Considering the overall importance of the BRCA2 BRC repeats in RAD51 binding and regulation, and the strong interactions we observed in our amylose pull-down approach, we decided to use BRC1-2 in our Y3H experiments. The Y3H approach was required for these experiments as previous work has shown individual RAD51 paralogs are not sufficiently expressed in Y2H assays without their cognate binding partners present to stabilize them[41, 76, 77]. Utilizing the Y3H assay, we observed interactions between BRC1-2 and RAD51B, RAD51C, RAD51D, XRCC2, and XRCC3, with RAD51B and RAD51D exhibiting the strongest interactions in the presence of endogenous yeast RAD51. We then used a Rad51Δ Y3H system to determine if any BRC1-2-RAD51 paralog interactions we observed were being mediated by yeast Rad51. Surprisingly, the yeast Rad51 mediated the BRC1-2-RAD51B interaction, as that interaction was lost in the Rad51Δ Y3H. The Y3H results not only confirm most of the interactions we observed in our amylose pulldown experiments, but also point towards these interactions being direct, at least for the BRC1-2 region of BRCA2 for RAD51D, XRCC2, and XRCC3. These results also solidify the RAD51B-RAD51 interaction, previously proposed by various labs[31, 32], by showing evidence that RAD51B-RAD51 interactions are evolutionarily conserved since yeast Rad51 can mediate an interact between human BRCA2 BRC1-2 and human RAD51B.

The BRC repeats of BRCA2 are known to be essential for the recruitment and stabilization of RAD51 nucleoprotein filament formation on ssDNA[13, 14, 18]. Direct visualization of RAD51 on ssDNA using negative-stain electron microscopy has shown that BCDX2 increases the nucleation and growth of RAD51 filaments[28, 31, 32]. Another report demonstrated that BCDX2 can dissociate from the RAD51 filament following formation and stabilization [32]. Taken together, our results suggest that the BCDX2 complex interacts directly with BRCA2, at least with BRC1-2 but likely other domains, and that RAD51B interacts with RAD51. These interactions likely reflect their ability to work conjointly to nucleate and elongate RAD51 nucleoprotein filaments. It is also likely the RAD51 paralogs have other direct interactions with BRCA2 that have yet to be characterized.

Although interactions were minimal to absent between RAD51C and BRCA2 in our amylose pull-down approach, the Y3H experiments were positive between BRC1-2 and RAD51C, albeit the weakest interactor amongst all the RAD51 paralogs. In the Rad51Δ Y3H, we saw loss of interaction between BRC1-2 and RAD51B where RAD51C was expressed as the RAD51B binding partner which means RAD51C was not able to bridge that interaction with BRC1-2. Though, in both the WT Y3H and Rad51Δ Y3H, there was growth in the BRC1-2 and RAD51C experiments. While these results initially seem to conflict, the Y3H was performed with the pRS416 vector expressing RAD51C’s binding partner, RAD51D. Considering that RAD51D displayed strong binding in both the amylose pull-down and the Y3H/Rad51Δ Y3H experiments, the interactions shown between BRC1-2 and RAD51C in the Y3H are likely mediated by RAD51D.

Our data indicate that the CTD of RAD51B, not the NTD, is responsible for the RAD51B-BRCA2 interaction. This finding is particularly interesting given the dynamic nature of the RAD51B CTD which prevented resolution in the cryo-EM structure. Disrupting the FxxA motif we identified in RAD51B with the F78A mutation abolishes the RAD51B interaction with BRCA2 but does not interfere with RAD51B-RAD51C complex formation further substantiating the importance of this interface in mediating the interaction with BRCA2. The results provide compelling evidence for a bona fide RAD51B-BRCA2 interaction and support the claim that F78 and A81 make up a previously unidentified FxxA motif that is crucial for RAD51B to bind BRCA2. Though, considering RAD51B-RAD51 interaction, further characterization of RAD51B(F78A)-RAD51 interaction is necessary to conclusively determine if RAD51B F78A is interrupting a direct interaction with BRCA2 or RAD51.

Knockout of individual RAD51 paralogs in human cells disrupts the formation of RAD51 foci in response to IR. Complementation with the respective wild-type RAD51 paralog rescues RAD51 foci formation. By complementing RAD51B knockout cells with either RAD51B-NTD or RAD51B-CTD, we determined that both the NTD and CTD of RAD51B are necessary for effective HR because neither domain alone could rescue RAD51 foci formation. While the partial rescue with the RAD51B F78A mutation demonstrates that a compromised FxxA motif decreases RAD51 foci formation efficiency, compensatory mechanisms must allow partial functioning. Since other RAD51 paralogs in the BCDX2 complex interact with BRCA2, RAD51D and XRCC2 could provide some compensation for the disruptions caused by F78A in RAD51B.

Our study deepens our understanding of BRCA2, RAD51, and the RAD51 paralogs by identifying and characterizing specific interactions between these critical proteins required for HR repair of DNA DSBs. We demonstrate direct interactions of both the BCDX2 and Cx3 complexes that take place at the BRC repeats of BRCA2, a key interaction hub allowing binding, loading, and stabilization of the RAD51 nucleoprotein filament. We further characterize RAD51B, whose CTD comprising the ATPase domain had previously been shown to regulate BCDX2 function, by determining that the CTD is responsible for mediating the BRCA2-RAD51B interaction. We also demonstrate that RAD51B interacts with yeast Rad51 giving strong evidence for RAD51B being the interface of BCDX2 that interacts with RAD51. We identified and validated a previously unrecognized FxxA motif in the linker region of RAD51B crucial for interaction with BRCA2. Using RAD51 foci formation in response to IR as an endpoint, we show that RAD51B functions are compromised in an F78A mutant further demonstrating the importance of the FxxA motif. Overall, this work brings fresh insights into new interactions between the RAD51 paralog complexes and BRCA2 which was previously unappreciated. We anticipate our findings will significantly impact clarification of how the RAD51 paralogs and BRCA2 work together to regulate RAD51 functions at DSBs and damaged replication forks.

## Materials and Methods

### 2.1 Cell culture

HEK293T cells and U2OS cells were cultured in DMEM with 10% fetal bovine serum (FBS). U2OS#18-RAD51B-8 (U2OS RAD51B KO) was cultured in DMEM with 15% FBS and 1xGlutaMAX supplement. U2OS RAD51B(NTD), U2OS RAD51B(CTD), U2OS RAD51B(WT), and U2OS RAD51B(F078A), were cultured in DMEM with 15% FBS, 1xGlutaMAX supplement, and 2 mg/mL G418. Transient transfections were carried out with Turbofect (Thermo Scientific R0531) following the manufacturer’s protocol. All cell lines were authenticated through ATCC STR profiling and tested regularly for mycoplasma using the Lonza MycoAlert detection kit (Lonza LT07-318).

### 2.2 Plasmid generation

Full-length wild-type human BRCA2 and BRCA2 domains (N-term, BRC1-8, DBD) were cloned into the phCMV1 mammalian expression vector with two tandem repeats of the maltose binding protein (2XMBP) tag located in-frame at the N-terminus of BRCA2 and the respective BRCA2 domains. An asparagine linker and a PreScission Protease cleavage sequence were placed between the 2XMBP tag and N-terminus of all proteins incorporated into this construct (previously described by Jensen et al., 2010)[13]. The BRCA2 was incorporated into the phCMV1 vector using a KpnI and NotI restriction digest and the BRCA2 domains were incorporated via a NotI and XhoI restriction digest strategy. 2XMBP N-term tagged RAD51B, RAD51C, RAD51D, XRCC2, and XRCC3 were also made in phCMV1 using the NotI and XhoI restriction digest strategy.

HA-RAD51B, HA-RAD51C, HA-RAD51D, HA-XRCC2, and HA-XRCC3 were cloned into the phCMV1 expression vector with the HA-tag located in-frame at the N-terminus of each paralog using a EcoRI and NotI restriction digest cloning strategy. RAD51B-HA-6xHis, RAD51C-HA-6xHis, RAD51D-HA-6xHis, XRCC2-HA-6xHis, and XRCC3-HA-6xHis was also cloned into the phCMV1 expression vector using an EcoRI and NotI restriction digest with an HA-tag and 6xHis-tag in-frame at the C-terminus of each paralog.

Takara PrimeStar Max (TakaraBio R047A) was used to PCR the inserts for cloning of 2XMBP-tagged and HA-tagged constructs following the manufacturer’s protocol. The RAD51B(F078A) mutagenesis was made using Q5 High-Fidelity 2X Master Mix (NEB M0492S) according to the manufacturer’s protocol.

### 2.3 Generation of stable cell lines

U2OS#18-RAD51B-8 (Leibniz-Institut DSMZ; originally generated by Garcin et al., 2019)[63] were stably transfected with 1 μg plasmid DNA using JetOptimus (Polyplus Transfection 101000006) following the manufacturer’s protocol in a 6-well plate. After 48 hours, the cells were trypsinized and diluted at ratios of 1:2, 1:4, 1:8, and 1:16 into 100 mm cell culture dishes with DMEM 15% FBS, 1xGlutaMAX supplement, and 2 mg/mL G418. After growing in selection, single cell colonies were picked into 96-well plates and then cultured into 24-well plates, 12-well plates, and 6-well plates. Protein expression was tested using western blotting and positive clones were selected and isolated for experimentation.

### 2.4 Western blots and amylose pulldowns

HEK293T cells were transfected in 6-well plates at 70% confluency using TurboFect reagent (Thermo Scientific) following the manufacturer’s protocol. 2XMBP tagged bait protein (Full-length BRCA2, BRCA2 domains, RAD51C, etc.) was co-expressed with HA-tagged prey protein (HA-tagged RAD51 paralogs). Untransfected HEK293Ts and the 2XMBP tag alone were used as negative controls. 48 hours after transfection the cells were lysed in 500 μL HEPES lysis bu]er (see bu]ers table) and treated with 0.5 μL Benzonase Nuclease (Millipore E8236). Total cellular lysate samples were taken for protein expression analysis, then the cell extracts were batch bound to washed amylose resin (NEB) for 2 hours to capture the 2XMBP-tagged proteins. The bound samples were washed 3 times with HEPES wash bu]er (see bu]ers table) then eluted with maltose elution bu]er (see bu]ers table). The total cell lysates and amylose pulldown samples were then run on a 4-15% gradient SDS-PAGE TGX stain-free gel (Bio-Rad), imaged using a ChemiDocMP imaging system (Bio-Rad), and then transferred to an Immobilon-P PVDF membrane (Millipore) in 1X Tris/glycine buffer (from 10X Tris/glycine buffer stock, Bio-Rad 161–0771). The membrane was blocked in 5% milk in 1X TBST. The washes were done using 1X TBST and the primary and secondary antibodies were diluted in 1X TBST for incubation. For the western blot, mouse antibodies against MBP (NEB E8032L, 1:20,000) were used to detect 2XMBP-tagged proteins and rabbit antibodies against HA (Cell Signaling C29F4, 3:20,000) were used to detect the HA-tagged RAD51 paralogs. After primary incubation, the blots were washed 3 times in 1X TBST and then incubated in secondary mouse and rabbit antibodies (m-IgGκ BP-HRP, Santa Cruz sc-516102, 1:10,000; mouse anti-Rabbit IgG-HRP, Santa Cruz sc-2357, 1:10,000, respectively). The western blots were developed using Clarity Western ECL substrate (Bio-Rad 170–5061) for 4 minutes, then imaged using the ChemiDocMP imaging system.

### 2.5 Immunofluorescence Imaging

RAD51B knockout U2OS cells and the stable cell lines expressing RAD51B(WT), RAD51B(NTD), RAD51B(CTD), and RAD51B(F78A) were grown on coverslips in 24-well plates. 20,000 cells were seeded on the coverslips and allowed to grow for 24 hours. Cells were either treated with 4 Gy ionizing radiation (IR) using an X-Rad 320 Biological Irradiator or left untreated as a control. Cells were fed with fresh media immediately after IR treatment. After 4 hours of treatment, cells were fixed and stained for immunofluorescence imaging. Cells were first washed twice with 1X PBS, fixed for 15 minutes using 1% paraformaldehyde 2% sucrose in 1X PBS at room temperature for 15 minutes, then washed twice again with 1 X PBS. The cells were then permeabilized for 30 minutes with methanol at −20°C, washed twice more with 1X PBS, then blocked with 4% BSA in PBS for 30 minutes at room temperature. After blocking the samples were incubated with primary antibodies against γH2AX (Millipore JBW301, 1:100) and RAD51 (Millipore Ab-1 PC130, 1:100) diluted in 4% BSA, 0.05% Triton X-100 in 1 X PBS, first for 60 minutes at 37°C then overnight at 4°C. The next day the cells were washed three times with 1X PBS, then incubated with secondary antibodies goat anti-rabbit Alexa Fluor 647 (Invitrogen A-21244, 1:1000) and goat anti-mouse Alexa Fluor 546 (A-11030, 1:1000) diluted in 4% BSA, 0.05% Triton X-100 in 1X PBS for 40 minutes at 37°C in a dark moist chamber. The coverslips were then washed with once with 1X PBS, incubated with 30 nM DAPI for 5 minutes, washed a final time with 1X PBS, then mounted on slides with FluorSave reagent (Millipore 345789). A Keyence BZ-X800E Fluorescent Microscope was used to take immunofluorescent images with a 60x objective lens.

### 2.6 Yeast-three-hybrid assays

For the yeast-three-hybrid (Y3H) experiment, the indicated *GAL4* activating domain (pGAD-RAD51B, RAD51C, RAD51D, XRCC2, XRCC3, RAD51, SWS1, empty), the *GAL4* DNA binding domain (pGBD-BRC1-2 and empty), and pRS416 vectors (pRS416-RAD51C, RAD51D, XRCC2, RAD51D, RAD51C, EMPTY, SWSAP1) were co-transformed into the *S. cerevisiae* competent PJ69-4α yeast strain[78]. The yeast were grown overnight in 5 mL of selective medium (SC-LWU) at 30°C to 0.4-0.6 OD_600_ as described previously[79]. Five µL of culture were spotted onto synthetic complete (SC) medium lacking leucine, tryptophan, and uracil to select for the plasmids as a loading control (SC-LWU) as well as medium lacking histidine (SC-HLWU), medium lacking histidine with 3AT [SC-HLWU+3-amino-1,2,4-triazole (3-AT)], and medium lacking adenine (SC-LWUA). Plates were grown for 2 days at 30°C and photographed. The images were adjusted identically for brightness and contrast using Adobe Photoshop.

### 2.7 Western blot analysis for Y3H experiments

PJ69-4α were transformed with either an empty pGBD plasmid or a pGBD plasmid expressing BRC1-2 with an HA tag. A single transformant was grown overnight in 5 mL of yeast extract peptone dextrose (YPD) at 30 °C. The cells were diluted to 0.2 OD_600_ in 5 mL YPD and grown for 3 h at 30°C. Whole-cell lysates of equal cell numbers (0.5 OD_600_) were prepared by TCA precipitation and 10 of 50 µL protein preparation was run on a 10% SDS-PAGE gel. The membrane was probed with HA antibodies (Abcam ab9110; 1:2500) to detect the HA-tagged protein, and Kar2 antibodies (Santa Cruz sc-33630; 1:200) as a loading control. Secondary antibodies used are anti-rabbit (1:10,000) The films were scanned and adjusted for contrast and brightness using Photoshop (Adobe Systems Incorporated).

### 2.8 Sequences and molecular modeling

Amino acid sequences for BRCA2, RAD51, and the RAD51 paralogs were obtained from the Uniprot Knowledgebase https://www.uniprot.org/ [80]. The UniProtKB codes of the sequence are: BRCA2 **P51587**, RAD51 **Q06609**, RAD51B **O15315-1**, RAD51C **O43502**, RAD51D **O75771**, XRCC2 **O43543**, and XRCC3 **O43542**.

The AlphaFold predicted structure of the BCDX2 complex was retrieved from ModelArchive https://www.modelarchive.org/ using accession code **ma-a54ps**, the Cryo-EM structure of the BCDX2 complex was retrieved from RCSB Protein Database[81] https://www.rcsb.org/ using the PDB ID **8OUY**[32] and the BRC4-RAD5 crystal structure was retrieved from RCSB PBD using PBD ID **1N0W**[66]

AlphaFold 3[82] was used to generate the predicted BCDX2 complex in Supplementary figure 4.2 using the following UniProtKB sequences: RAD51B **O15315-1**, RAD51C **O43502**, RAD51D **O75771**, XRCC2 **O43543**, and XRCC3 **O43542**. The predicted structure has an ipTM = 0.8 and a pTM =0.82.

PyMOL Molecular Graphics System, version 2.5.0, Schrödinger, LLC, was used for visualization of the AlphaFold predicted models and Cryo-EM molecular structures of the BCDX2 complex.

## Acknowledgements

We thank all members of the Jensen lab for their insights, guidance, and discussion. We would like to thank Dr. Gemma Moore for her input and feedback. This research was supported by a grant from the NIH (R21 ES034164-01), and J.G.T was also supported by a training grant and fellowship from the NIH (T32 GM007223, F31 CA265150-03). K.A.B. is supported by NIH (R01 ES030335, R01 ES031796) and Department of Defense (BC201356).

---

**Supplementary Figure 1.**
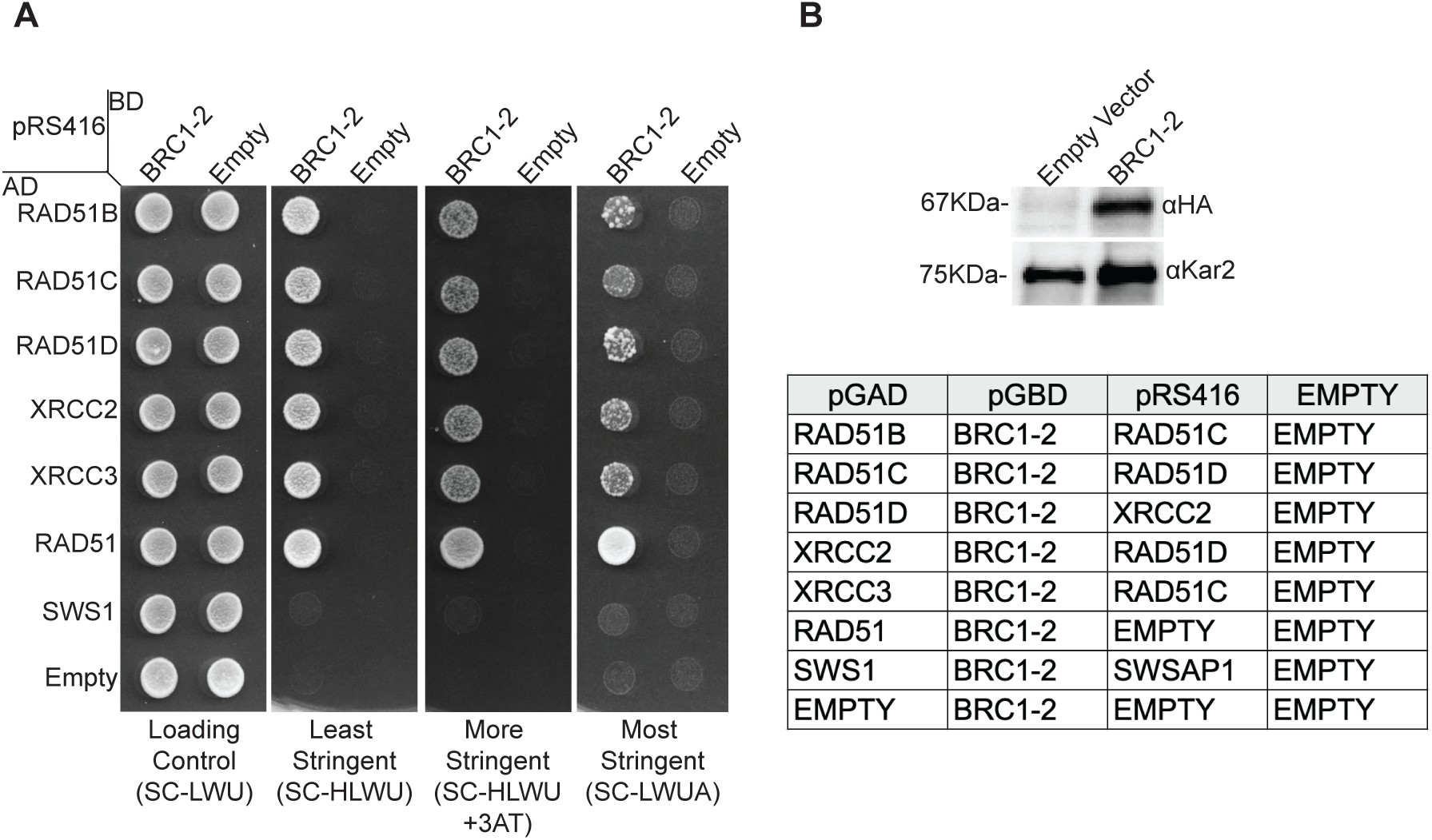
BRC1-2 repeats exhibit a Y3H interaction with members of BCDX2 and CX3 complexes. **(A)** Yeast-three-hybrid (Y3H) analysis of HA-tagged BRC1–2 with RAD51 paralogs. Cells expressing GAL4 BD–BRC1–2 or empty vector, GAL4 AD–RAD51 paralogs or empty vector, and a pRS416 vector stabilizing RAD51 paralogs with their binding partners were assayed for interaction. Equal cell numbers were spotted on control and selection plates. Growth after 2 days at 30°C indicates a Y3H interaction. Experiments were performed in triplicate; representative images are shown. **(B)** Protein expression of BRC1–2 in yeast was confirmed by Western blot. Whole-cell lysates expressing empty vector or HA-tagged BRC1–2 were probed with anti-HA and anti-Kar2 antibodies. Kar2 served as a loading control.

**Supplementary Figure 2.**
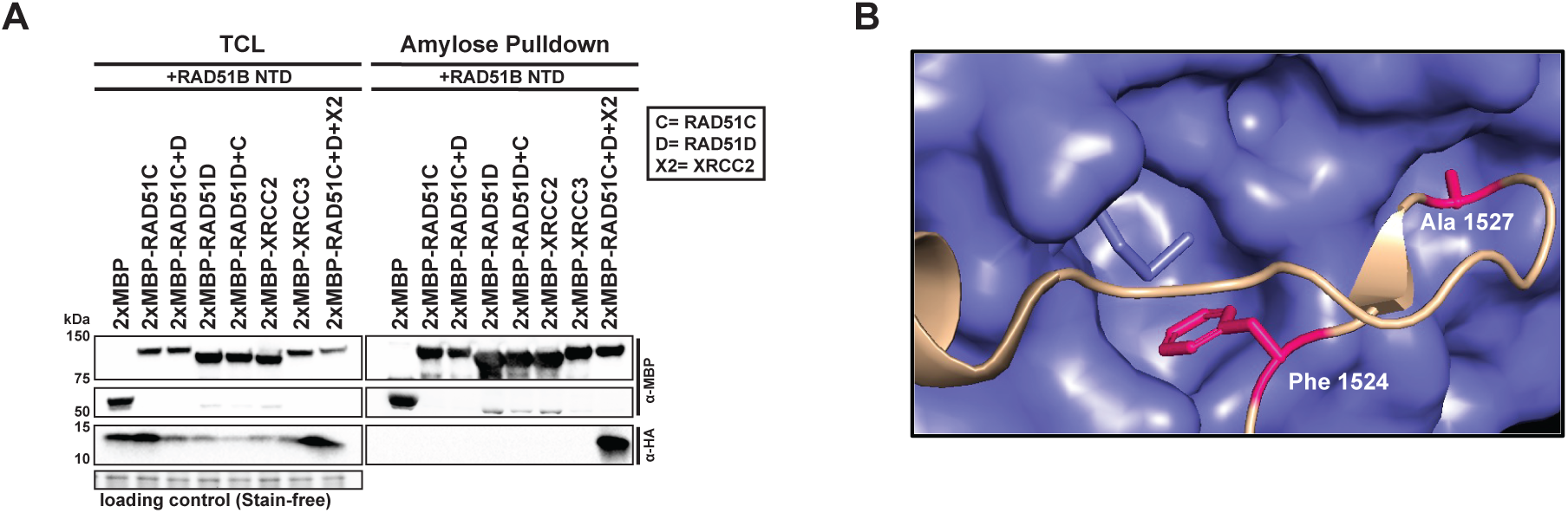
RAD51B NTD requires other BCDX2 members for NTD stacking interaction; FxxA motif mediates BRC4–RAD51 interaction. **(A)** RAD51B NTD requires all BCDX2 components to stabilize NTD stacking. Western blots of amylose pulldowns showing that RAD51B NTD is pulled down by 2XMBP-RAD51C only when RAD51D and XRCC2 are co-expressed. **(B)** FxxA motif mediates BRC4–RAD51 interaction. Crystal structure [PDB: 1N0W] of BRC peptide (light brown) bound to RAD51 (purple), highlighting the FxxA motif (pink). The phenylalanine and alanine residues interact with hydrophobic pockets on RAD51.

**Supplementary Figure 3.**
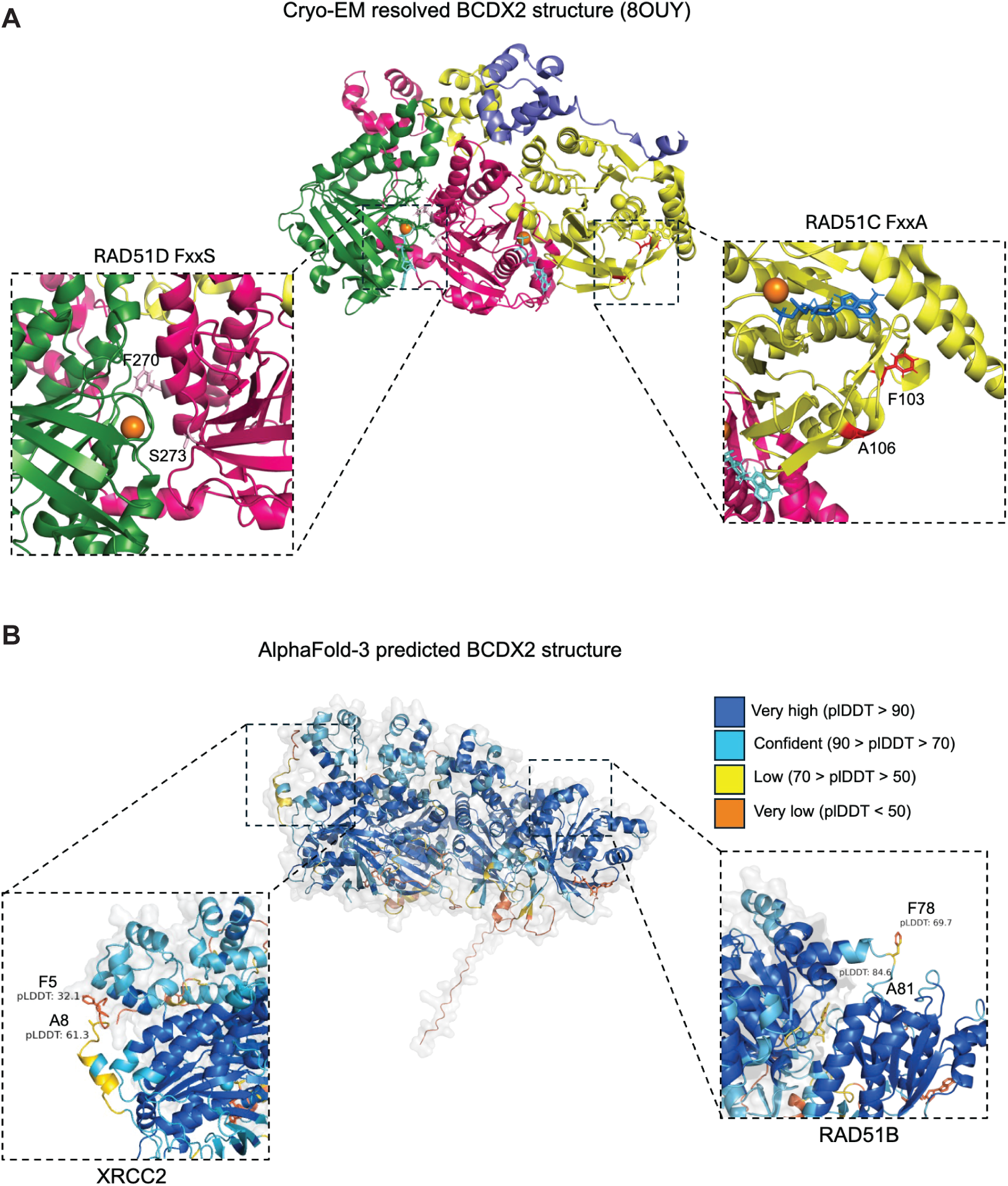
Location of FxxA and FxxS motifs in BCDX2 complex structures. **(A)** Cryo-EM structure of BCDX2 highlighting RAD51D FxxS and RAD51C FxxA motifs. The complex includes RAD51B NTD (purple), RAD51C (yellow), RAD51D (pink), and XRCC2 (green). Mg²⁺ ions (orange spheres), ATP (light blue), and ADP (dark blue) are shown. Insets provide zoomed views of RAD51D FxxS (light pink) and RAD51C FxxA (red) motifs. **(B)** AlphaFold-predicted BCDX2 structure highlighting RAD51B FxxA and XRCC2 FxxA motifs. The structure is color-coded by pLDDT confidence scores. Insets zoom in on XRCC2 FxxA and RAD51B FxxA motifs, with pLDDT values for key residues included.

